# *Candida albicans* hyphal expansion causes phagosomal membrane damage and luminal alkalinization

**DOI:** 10.1101/340315

**Authors:** Johannes Westman, Gary Moran, Selene Mogavero, Bernhard Hube, Sergio Grinstein

**Author notes:** These authors contributed equally (Gary Moran, Selene Mogavero). Corresponding author: Sergio Grinstein.

## Abstract

Macrophages rely on phagosomal acidity to destroy engulfed microorganisms. To survive this hostile response, opportunistic fungi such as *Candida albicans* developed strategies to evade the acidic environment. *C. albicans* is polymorphic, able to convert from yeast to hyphae, and this transition is required to subvert the microbicidal activity of the phagosome. However, the phagosomal lumen, which is acidic and nutrient-deprived, inhibits yeast-to-hypha transition. To account for this apparent paradox, it was recently proposed that *C. albicans* produces ammonia that alkalinizes the phagosome, thus facilitating yeast-to-hypha transformation. We re-examined the mechanism underlying phagosomal alkalinization by applying dual-wavelength ratiometric pH measurements. The phagosomal membrane was found to be highly permeable to ammonia, which is therefore unlikely to account for the pH elevation. Instead, we find that yeast-to-hypha transition begins within acidic phagosomes, and that alkalinization is a consequence of proton leakage induced by excessive membrane distension caused by the expanding hypha.

**IMPORTANCE:** *C. albicans* is the most common nosocomial fungal infection, and over three million people acquire life-threatening invasive fungal infections every year. Even if antifungal drugs exist, almost half of these patients will die. Despite this, fungi remain underestimated as pathogens. Our study uses quantitative biophysical approaches to demonstrate that the yeast-to-hypha transition occurs within the nutrient deprived, acidic phagosome and that alkalinization is a consequence, as opposed to the cause of hyphal growth.

## INTRODUCTION

*Candida albicans* is a commensal yeast of humans, but is frequently the source of mucosal infections and can, in severe cases, cause life-threatening systemic infections (1). It colonizes the epithelial surfaces of 30-70% of healthy individuals, and superficial infections are usually transient (2). Due to an aging population, an increased use of antibiotics and immunocompromising drug treatments, nosocomial *C. albicans* infections have increased dramatically over the last decades (3). Unlike many other pathogenic microbes, *C. albicans* is polymorphic and grows as budding yeast, pseudohyphae or true filamentous hyphae. The yeast-to-hypha transition is initiated as a response to various environmental stimuli. These include increased pH or temperature, nutrient deprivation, contact with immune cells, and exposure to serum proteins (4, 5). *C. albicans* yeast cells are associated with commensal growth (but also with dissemination via the blood stream); by contrast, hyphae are capable of invading epithelia, endothelia and organ tissues, and are thus essential for pathogenicity (4, 5).

*C. albicans* colonization of the gut is restricted by the bacterial microbiota, but also by the immune system, notably patrolling phagocytes. To prevent microbial dissemination from the gut, macrophages and neutrophils quickly recognize and engulf invading microbes through phagocytosis. After engulfment of microbial cells by macrophages, the nascent phagosome undergoes a series of fusion and fission events with endosomes and lysosomes, a phenomenon referred to collectively as ‘phagosome maturation’ (see Walpole, Grinstein and Westman, 2018 for a review). Fusion of the nascent phagosome with endosomes and lysosomes induces a progressive luminal acidification. This is attributed to the gradual acquisition of vacuolar proton ATPases (V-ATPases) from endosomal compartments. The prevailing phagosomal pH dictates the efficiency of microbial killing and antigen presentation, as well as the degradation of the ingested prey (7).

After phagocytosis of *C. albicans* yeast, the fungus is confined within the mature phagosome. Nonetheless, at least *in vitro, C. albicans* can escape as a result of intra-phagosomal hyphal formation (8–10). It is currently believed that the yeast-to-hypha transition is inhibited within acidic phagosomes, and, consequently, *C. albicans* is thought to manipulate the phagosomal pH prior to hypha formation (11–18). Thus, the ability of *C. albicans* to alkalinize the phagosome is considered crucial for survival and escape from the macrophage.

Recent studies have proposed that phagosomal alkalinization is a consequence of ammonia (NH_3_) release by *C. albicans* (15–18). NH_3_ can in principle alkalinize the phagosome by consumption of protons and formation of ammonium (NH_4_^+^). It has been further proposed that, after sufficient NH_3_ production and associated proton consumption, hyphal formation can occur, followed by eventual escape from the phagosome and ultimately from the macrophage itself. Besides *C. albicans*, NH_3_ generation and protonation has also been suggested to be the cause of phagosome alkalinization for other pathogens such as *Mycobacterium tuberculosis* and *Helicobacter pylori* (19–22).

To effectively mediate phagosomal alkalinization, NH_3_ production has to exceed the rate of proton pumping by the V-ATPases. Moreover, and most importantly, the rate of NH_3_ generation has to exceed the rate at which NH_3_ diffuses out of the phagosome. In this regard, it is noteworthy that most mammalian membranes are highly permeable to NH_3_ (23–27). Validation of the alkalinizing role of NH_3_ therefore requires quantitative comparison of these parameters.

A second mechanism that could affect phagosomal pH, which is not mutually exclusive with the generation of NH_3_, is proton exit from the phagosome via Candidalysin (28). This pore-forming toxin is a hydrophobic, α-helical peptide secreted by *C. albicans* hypha after cleavage of the polyprotein Ece1 (28, 29). Candidalysin has been shown to intercalate into membranes and to form pores, leading to lysis of epithelial cells. However, its role during interaction with macrophages and its potential ability to permeabilize the phagosomal membrane has not been investigated.

In this study, we analyzed the role of NH_3_ in phagosome alkalinization by *C. albicans*. By applying dual-wavelength ratiometric fluorescence imaging, we undertook measurements of phagosomal buffering power, rate of proton pumping and phagosomal NH_3_ permeability, and compared them to the rate of NH_3_ production by *C. albicans*. We also investigated the role of Candidalysin, and assessed whether hyphal growth itself contributed to phagosomal alkalinization.

## RESULTS

### The rate of proton pumping by V-ATPase surpasses the rate of NH_3_ production by *C. albicans*

For NH_3_ generation by *C. albicans* to account for macrophage phagosome alkalinization, it would need to exceed the rate of proton pumping by the phagosomal V-ATPases (Figure 1A). We calculated proton pumping by measuring the rate of change of pH (ΔpH/Δt) induced by addition of the potent and specific inhibitor concanamycin A (CCA) to the murine macrophage cell line RAW264.7 (hereafter referred to as RAW cells). Such measurements are based on the notion that, in the steady state, the rate of pumping by the V-ATPases is identical to the rate of proton (equivalent) leakage (30). To measure the phagosomal pH, *C. albicans* yeast cells were allowed to bind FITC-labeled concanavalin A and a *C. albicans-specific* IgG prior to phagocytosis. Such labeled and opsonized yeast cells were centrifuged onto macrophages grown on glass coverslips to initiate phagocytosis synchronously and, at the desired times, phagosomal pH was measured by dual-wavelength ratiometric fluorescence imaging as detailed under Methods.

**Figure 1.**
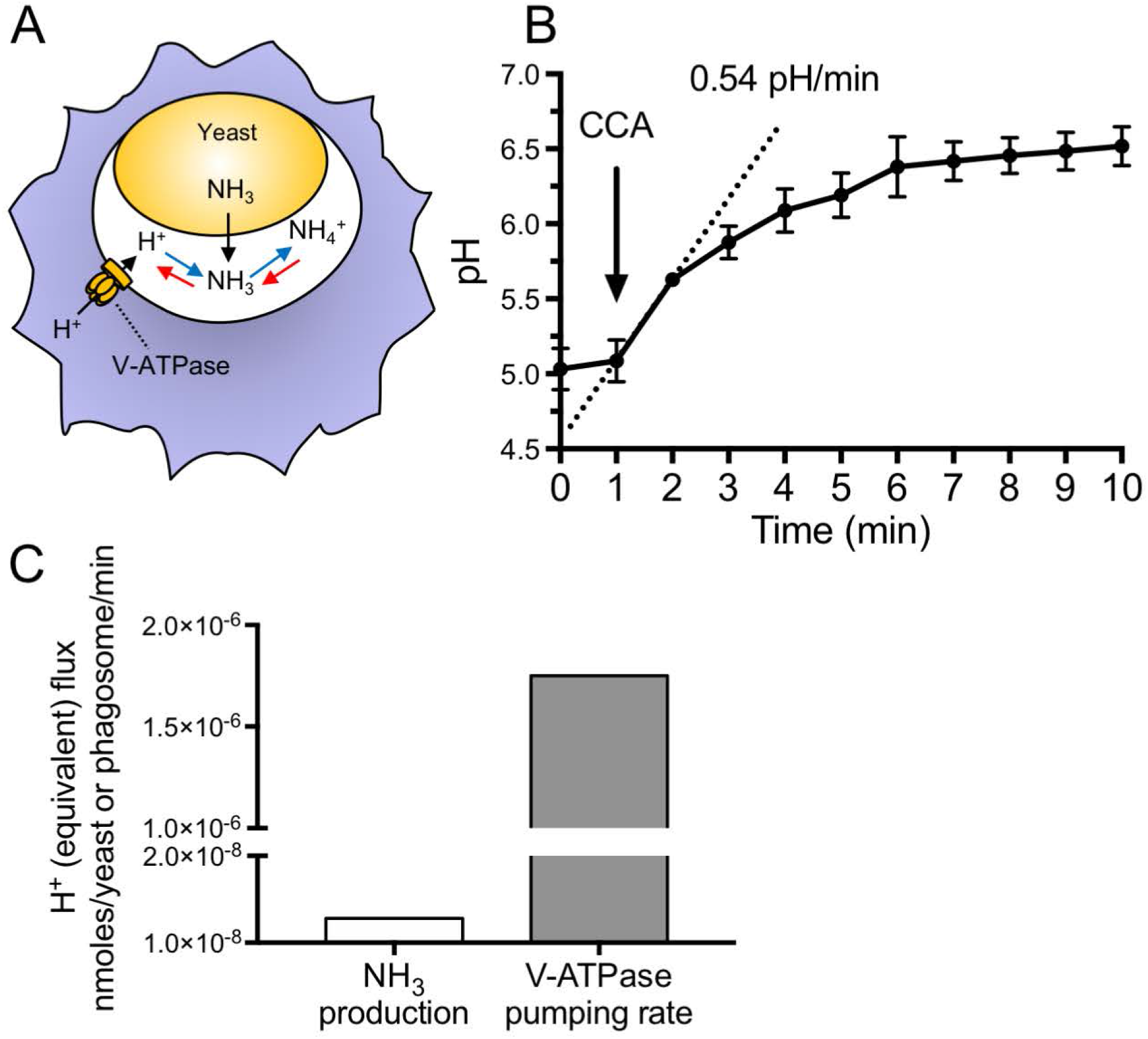
The rate of proton pumping by the V-ATPase exceeds the rate of NH_3_ production. (A) Schematic model illustrating proton pumping by the V-ATPase, NH_3_ production by the yeast and its protonation to NH_4_^+^. (B) Phagosomal pH measurements acquired by fluorescence ratio imaging. After recording the baseline pH, Concanamycin A was added where indicated and pH was measured for a further 9 min. The initial rate of ΔpH, estimated from the slope of the dotted line, is shown. Data are means ± SEM of 16 determinations in 4 independent experiments. (C) Comparison of the rates of NH_3_ production and V-ATPase pumping. Proton pumping was calculated from experiments like that in B, as described in Materials and methods, while the rate of NH_3_ production was derived from (15).

As illustrated in Figure 1B, when added 1 h after phagocytosis –when the phagosomes are acidic, averaging a pH of 5.03 ± 0.14 (mean ± SEM of 32 determinations in 4 experiments)– CCA elicited a rapid alkalinization at an average rate of 0.54 pH/min. The amount of protons pumped per unit time can be calculated by multiplying this rate times the phagosomal buffering capacity. The latter was measured by pulsing the cells with known amounts of membrane-permeable weak electrolytes (see Methods and Loiselle and Casey, 2010). In 4 independent experiments, the phagosomal buffering power averaged 91.2 ± 3.3 mmoles/L/pH. The rate of proton pumping at the steady state was therefore calculated to be 49.2 ± 15.5 mmoles/L/min.

We proceeded to compare the rate of pumping with the reported rate of NH_3_ production by *C. albicans*. Vylkova and Lorenz (15) reported a production of ≈35 ppm over 24 hrs, which is equivalent to 1.28 * 10^−8^ nmoles/yeast/min. Similar rates have been reported by others (13, 14, 16–18). This rate is two orders of magnitude lower than the rate of proton pumping at the steady state (Figure 1C). It should be noted that the activity of the V-ATPase decreases markedly as the pH becomes more acidic (32), so that the disparity between the rates of pumping and NH_3_ production would become even greater at more alkaline pH. At such pH values the rate of leakage of proton equivalents by other (endogenous) pathways decreases, which would further offset the rates of acidification and alkalinization. We conclude that NH_3_ production by *C. albicans* is unlikely to account for the reported phagosomal alkalinization.

### *C. albicans-containing* phagosomes are permeable to NH_3_

Not only is the rate of NH_3_ production insufficient to overcome the rate of proton pumping, but sustained alkalinization would require retention of NH_3_ within the phagosome. Because, as illustrated diagrammatically in Figure 2A, the protonation of NH_3_ is a rapidly reversible reaction, a fraction of the NH_3_/NH_4_^+^ will always exist inside phagosomes as the unprotonated weak base. Because the extracellular space and the cytoplasm are nominally free of NH_3_/NH_4_^+^, the prevailing outward gradient would promote ongoing loss of NH_3_ from the phagosome, provided the phagosomal membrane is permeable to the weak base. While most mammalian membranes are permeable to NH_3_, the permeability of the *C*. albicans-containing phagosome has not been ascertained. We assessed NH_3_ permeability by measuring the phagosomal pH while pulsing the medium with extracellular NH_3_/NH_4_^+^, as illustrated in Figure 2B. Addition of NH_3_/NH_4_^+^ to the medium caused an immediate and pronounced phagosomal alkalinization (Figures 2C-E), ostensibly due to permeation of NH3 and protonation to NH_4_^+^. Of note, acute removal of extracellular NH_3_/NH_4_^+^ resulted in rapid restoration of the acidic pH, implying very rapid conversion of NH_4_^+^ to NH_3_ and efflux of the latter. Such experiments enabled us to estimate the rate at which NH_3_ permeates the membrane of *C. albicans*-containing phagosomes. Considering the ΔpH elicited by NH_3_/NH_4_^+^ (from 4.85 ± 0.22 to 6.0 ± 0.03, n= 4; Figure 2E) after 1 sec –the fastest time measurable using our experimental set-up– and the buffering power determined earlier, we estimated that NH_3_ can enter/exit phagosomes at a rate of 2.42 * 10^−4^ nmoles/phagosome/min. As illustrated graphically in Figure 2F, this rate is several orders of magnitude greater than the reported rate of NH_3_ production by *C. albicans*. These additional data reinforce our conclusion that NH_3_ production by *C. albicans* is unlikely to account for the reported phagosomal alkalinization.

**Figure 2.**
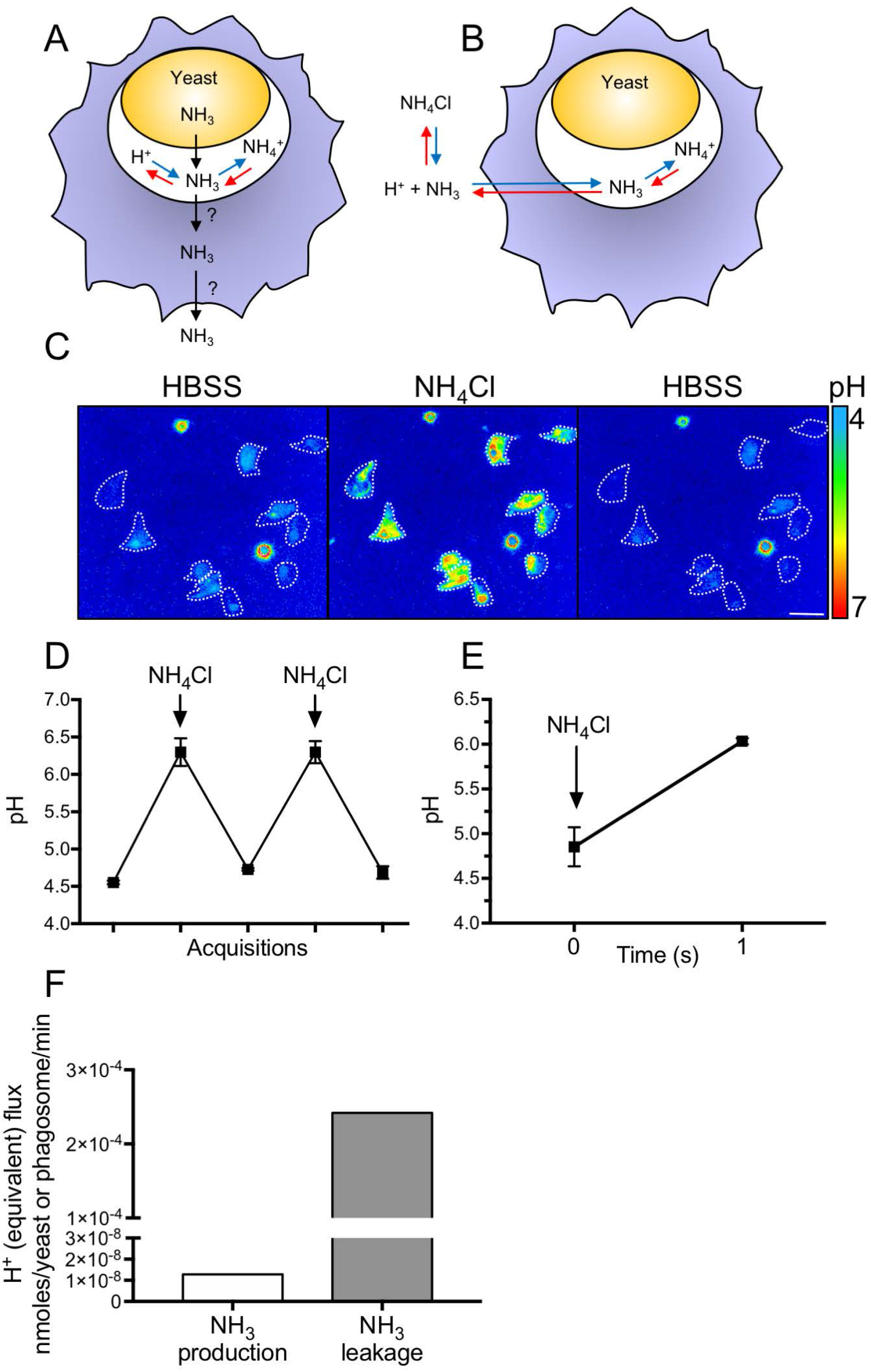
Phagosomes containing *C. albicans* are highly permeable to NH_3_. (A) Schematic model illustrating *C. albicans* production and secretion of NH_3_, which can be protonated to form NH_4_^+^. The possibility that NH_3_ diffuses out of the phagosome and the cell is considered. (B) Schematic model illustrating the possible fate of NH_3_ following addition of NH_4_Cl to the extracellular milieu (blue arrows) and its subsequent removal (red arrows). (C) Representative pseudocolor images showing the phagosomal pH before (left), during (middle) and after (right) bathing cells in medium containing 15 mM NH_4_Cl. Scale bar = 15 μm (D) Measurement of phagosomal pH during repeated exposure to and removal of extracellular NH_4_Cl. Data are means ± SEM of 24 determinations in 4 independent experiments. (E) The phagosomal pH was measured before and 1 s after addition of 15 mM NH_4_Cl. Data are means ± SEM of at least 15 determinations in 4 independent experiments for each type. (F) Comparison of the rates of NH_3_ production and NH_3_ leakage. The latter was calculated from experiments like those illustrated in D, as described in Materials and methods.

### *C. albicans* hyphal expansion drives phagosomal alkalinization

We next analyzed the time course of the pH changes undergone by the *C*. albicans-containing phagosomes (Figure 3A). While NH_3_ is presumably produced continuously by the yeast, the phagosome initially becomes acidic and remains so for nearly 2 hrs. These observations not only argue once again against a role for NH_3_ production in the pH changes, but suggest that an alternative, time-dependent mechanism is involved. Significant alkalinization was detectable only after ≈3 hrs. Notably, marked hyphal growth was clearly apparent at this stage: on average the hyphae reached 24.7 ± 2 μm in length (Figure 3A). We therefore hypothesized that hypha can form inside acidic phagosomes, and that phagosomal alkalinization was a consequence of hyphal growth.

**Figure 3.**
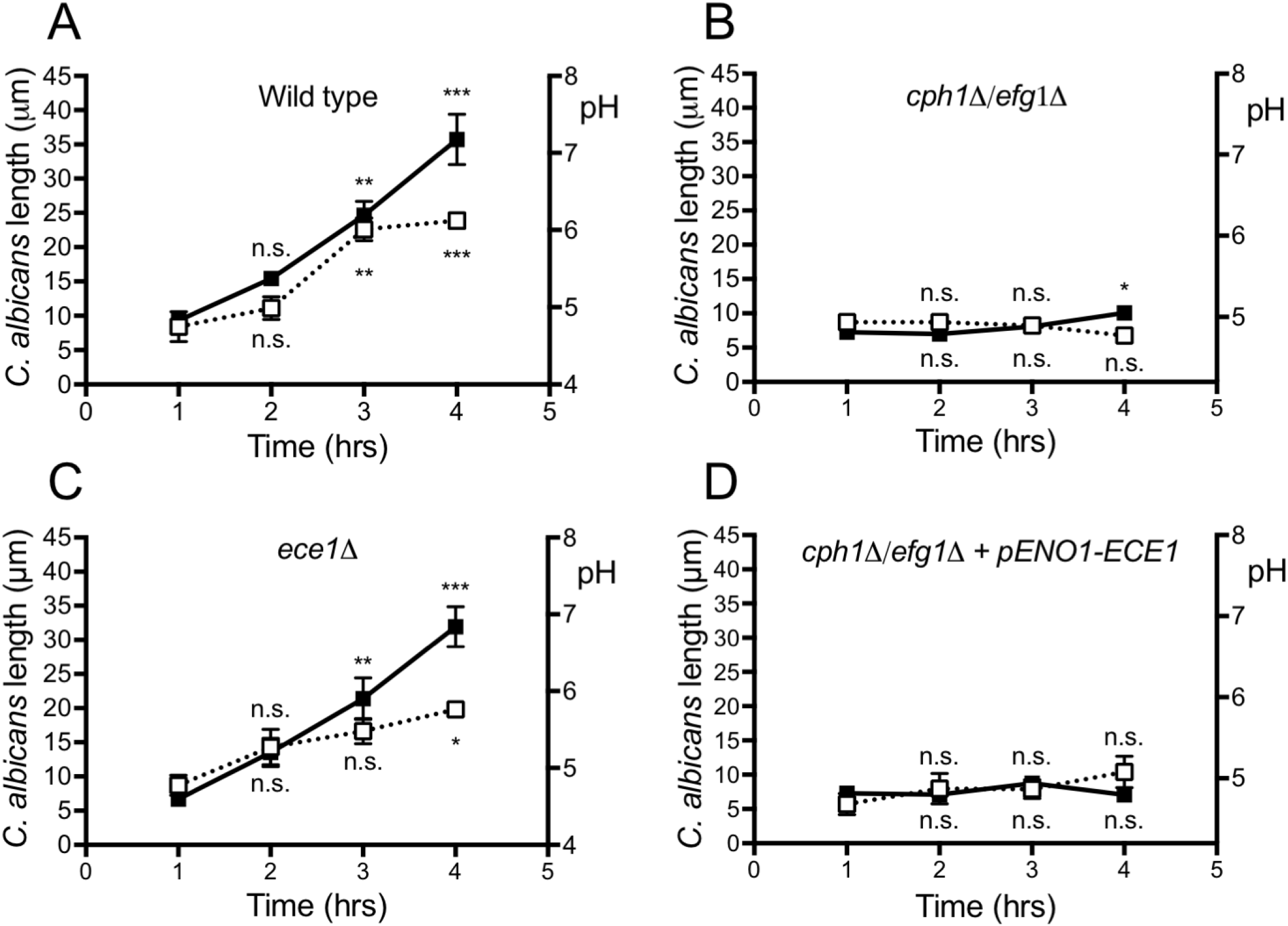
Phagosome alkalinization is associated with hyphal extension. (A-D) Phagosomal pH (right y-axis) and maximal length (left y-axis) were measured as a function of time following phagocytosis of the indicated *C. albicans* strains (see Methods for details). In each instance, at least 30 phagosomes were quantified per time point and at least 3 independent experiment were performed per condition. Data show means ± SEM. Significance was calculated by one-way ANOVA, with Tukey’s test. * indicates p ≤ 0.05, ** p ≤ 0.01, and *** p ≤ 0.001.

To test this hypothesis, we investigated whether the yeast-locked *cph1*Δ/*efg1*Δ (*C. albicans* mutant (33), which is unable to form hyphae, alters the phagosomal pH (as does the wild type. Remarkably, over a period of four hours yeast-locked *C. albicans* cells failed to dissipate the phagosomal acidification (Figure 3B), despite the fact that this strain generates NH_3_ at rates comparable to the wild type *C. albicans* (Figure A2A). These data are consistent with the notion that the pH change is a consequence of hyphal growth.

During hypha formation, the expression of several genes is activated (4). One such gene is *ECE1*, the product of which is processed into the pore-forming toxin Candidalysin (28). Because of the association between hyphal growth and phagosomal pH changes, we tested whether Candidalysin contributes to the alkalinization. The pores formed by the toxin could conceivably cause leakage of proton (equivalents) through the phagosomal membrane. As illustrated in Figure 3C, a mutant lacking Ece1 (*ece1*Δ), the precursor required for Candidalysin generation, caused phagosomal alkalinization at a rate that was only slightly slower than that induced by the wild type. The difference between the two strains was small but statistically significant (Figure A2B). Of note, the *ece1*Δ mutant formed hyphae that grew inside the phagosome at rates that were similar to the wildtype (Figure AS2C).

These results indicate that, while Candidalysin aids in alkalinizing the phagosome, its contribution is comparatively small and that other factors are involved. To validate this conclusion, we constructed a strain of yeast-locked *C. albicans* that expresses *ECE1* at levels comparable to those recorded during hyphal growth of wild type *C. albicans* and that produced similar quantities of the Candidalysin peptide (Data-set A1 and Figure A1). Despite the continuous production of Candidalysin, this strain had negligible effects on phagosomal pH over a four-hour period (Figure 3D).

The results described above suggest that the phagosomal pH change is a direct consequence of hyphal growth. Remarkably, the opposite has previously been proposed: namely that alkalinization precedes and is required for hyphal growth (15, 34). The latter concept derives from analyses of the pH dependence of *C. albicans* hyphal growth, which show faster growth rates at more alkaline pH (34–37). Indeed, it has been shown that neutral pH is required for full virulence of *C. albicans*, as many effectors are activated by proteolytic cleavage at neutral pH (38). However, while we could readily replicate the faster growth of *C. albicans* at more alkaline pH values, we also noted that hyphal growth does occur *in vitro* at the pH normally attained by phagosomes, i.e. pH 4.5-5.0 (Figures 1–3 and Figure A2).

### *C. albicans* hyphal expansion drives phagosomal membrane rupture

As the hypha expands over time within the phagosome, increasing mechanical tension must be applied on the phagosomal membrane. This could conceivably alter the permeability of the membrane to proton (equivalents) and potentially even cause its rupture.

We assessed the phagosomal membrane integrity using sulforhodamine B (SRB), a fluorescent dye that is nominally impermeable across biological membranes. SRB was delivered to phagosomes via fusion with lysosomes, which had been previously loaded with the dye using a pulse-chase protocol (see Methods for details). As illustrated in Figure 4A and Movie A1, SRB contained within phagolysosomes can be initially observed lining the yeast and hyphae. Over time, however, the SRB contained within phagosomes formed by wild type *C. albicans* was lost progressively (Figures 4A-B). In line with phagosomal alkalinization, the number of SRB-positive phagosomes decreased over a period of 4 hrs (Figure 4B). SRB was lost at a similar rate from the *ece1*Δ mutant, which grows at a comparable rate to the wild type *C. albicans* strain (Figure 4C). Importantly, SRB was retained during the same period by the yeast-locked mutants, even when they were engineered to produce Ece1 (Figures 4D-E). The latter data suggest that SRB-leakage is a consequence of hyphal growth, and not permeation of the dye via Candidalysin. Our data are most consistent with a growth-induced change in permeability, likely manifested as rupture of the membrane. Note that, as documented below, in some instances rupture may have been transient, followed by membrane repair.

**Figure 4.**
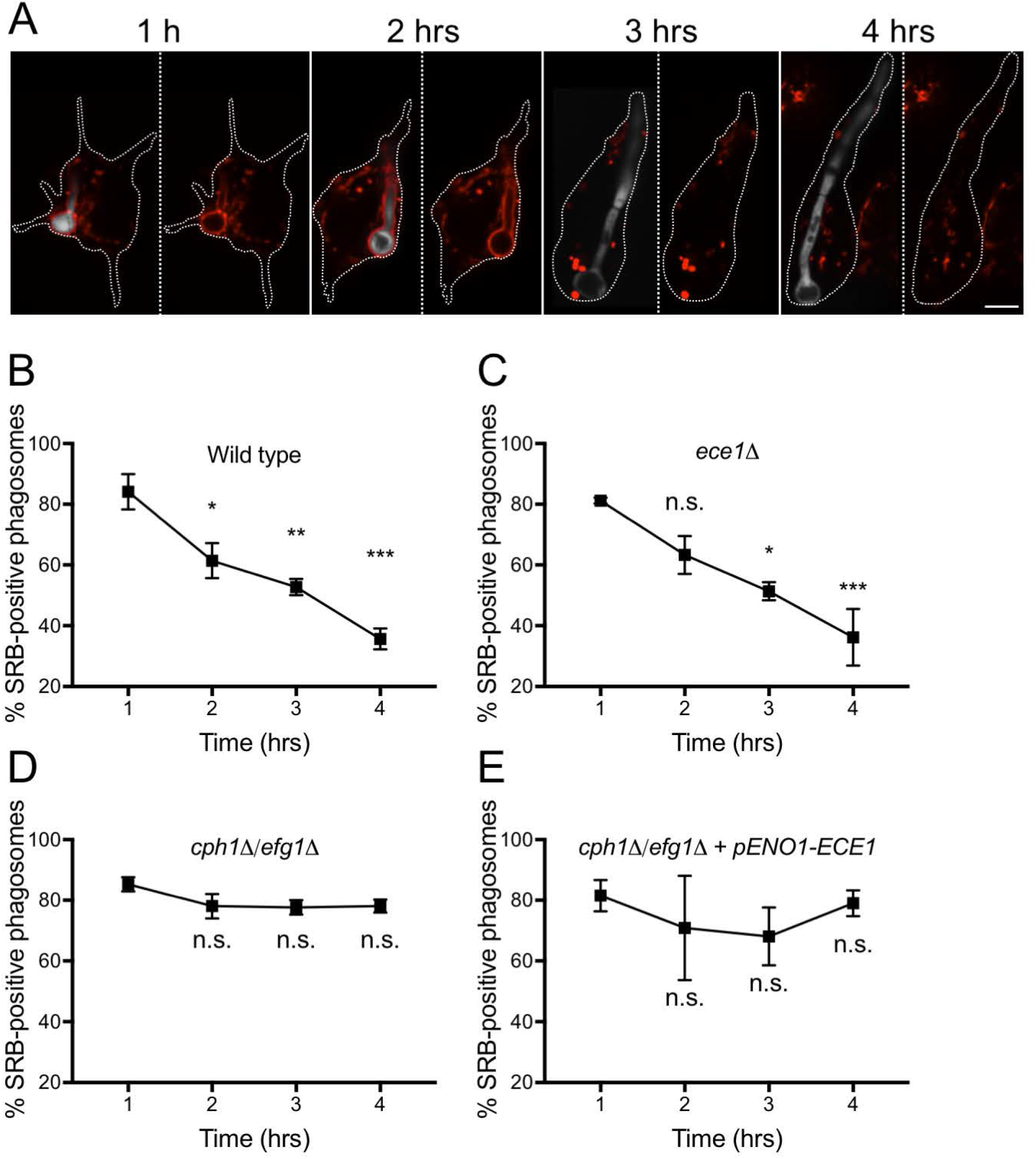
Growth-induced phagosomal leakage demonstrated using Sulforhodamine B (SRB). To load endosomes and lysosomes with SRB (red), cells were bathed in 150 μg/mL of the dye 60 min prior to phagocytosis of wild-type *C. albicans* yeast. (A) During phagosome maturation, endosomes and lysosomes fuse with the phagosome, delivering SRB to its lumen. Cells were imaged over 4 hrs following phagocytosis by spinning-disk confocal microscopy and representative images are depicted. *C. albicans* is shown in white (left images for each time); the yeast were omitted from the right panels to more clearly visualize the SRB outline. The borders of the macrophages are outlined using a dotted white line. Scale bar = 5 μm. (B-E) Quantitation of the fraction of phagosomes retaining SRB as a function of time after phagocytosis. Results obtained using wild-type *C. albicans* (B) or Ece1-null (*ece1*Δ) (C), yeast locked (*cph1*Δ/*efg1*Δ) (D) or Ece1-expressing yeast-locked (*cph1*Δ/*efg1*Δ + *pENO1-ECE1*) (E) strains are illustrated. Data are means ± SEM of at least 100 determinations in 3 independent experiments for each type. Significance was calculated using one-way ANOVA, with Tukey’s test. * indicates p ≤ 0.05, ** p ≤ 0.01, and *** p ≤ 0.001. n.s.= not significantly different.

### Phagosomal expansion using GPN causes alkalinization and membrane rupture similar to hyphal expansion

To verify that mechanical tension (like that induced by hyphal growth) suffices to produce phagosomal alkalinization and membrane rupture, we treated phagosomes containing the yeast-locked *C. albicans* mutant *cph1*Δ/*efg1*Δ with Gly-Phe-β-napthylamide (GPN). GPN is a membrane-permeable dipeptide and a substrate for cathepsin C. Phagosomal cathepsin C can cleave GPN, generating the membrane-impermeable Phe-β-napthylamide. Accumulation of Phe-β-napthylamide drives osmotically obliged water into the phagosome, which consequently expands, exerting hydrostatic pressure that distends the membrane in a manner akin to that occurring during hyphal growth. As illustrated in Figure 5A (left panel and Movie A2), following addition of GPN, phagosomes containing the yeast-locked *C. albicans* expanded rapidly and alkalinized within 20 min, while untreated phagosomes remained acidic (Figure 5A, right panel and Movie A2). Even faster onset of alkalinization was observed using higher concentrations of GPN (not shown). GPN also induced leakage of phagosomal SRB within minutes (Figure 5B, left panel and Movie A3), while untreated phagosomes remained SRB-positive (Figure 5B, right panel and Movie A3). We concluded that mechanical tension is sufficient to cause phagosome alkalinization and membrane rupture.

**Figure 5.**
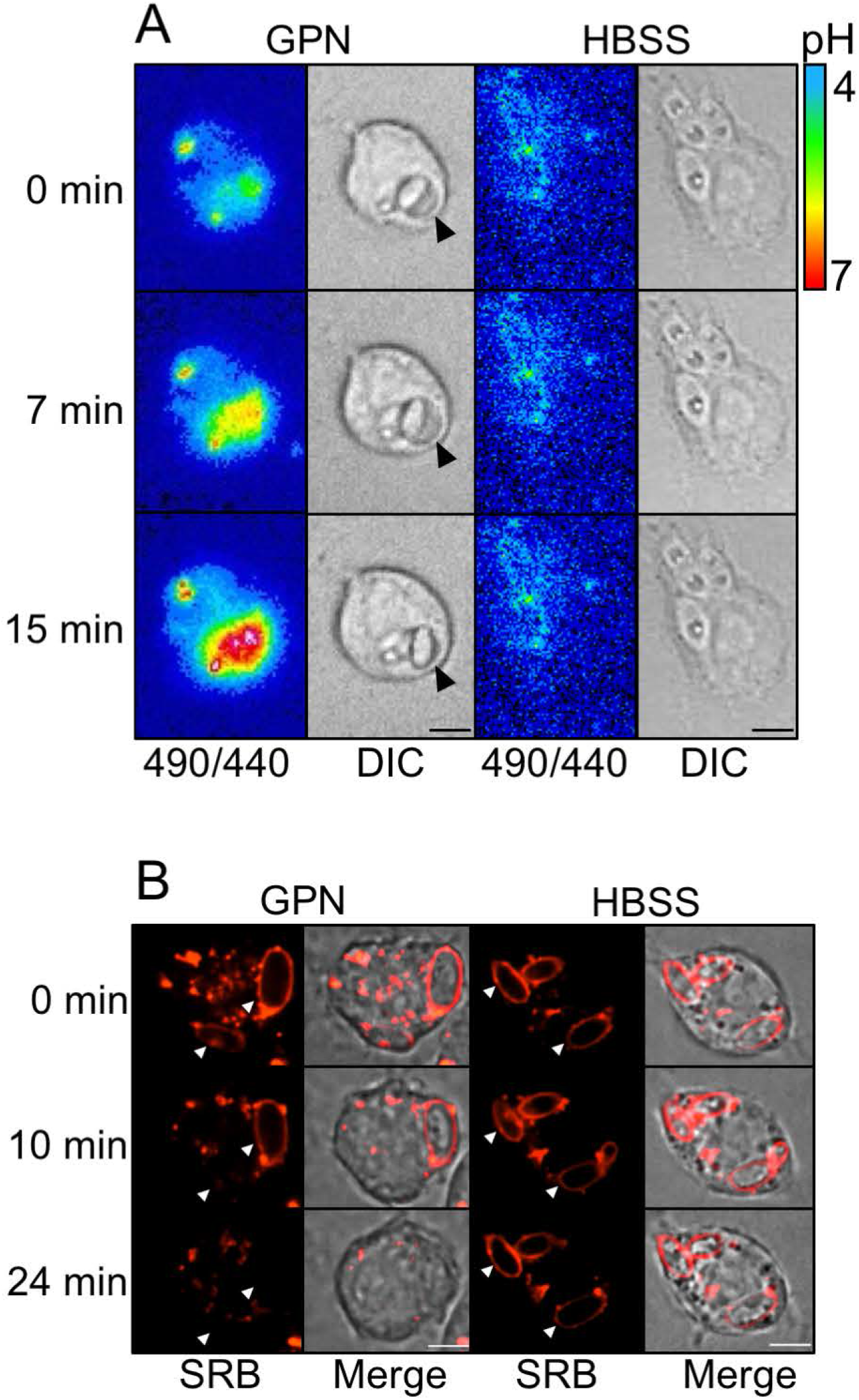
GPN induces expansion-dependent phagosomal alkalinization and membrane rupture. (A) One hour after phagocytosis of yeast-locked *C. albicans*, phagosomal expansion was induced by adding 100 μM GPN. Phagosomal pH was recorded every 15 seconds by fluorescence imaging, measuring the ratio of the emission obtained when exciting at 490 nm vs. 440 nm. Vehicle alone (HBSS) was used as negative control. Scale bar = 5 μm. (B) Endosomes and lysosomes were loaded with SRB (red) by incubating cells with 150 μg/mL of the dye for 60 min prior to phagocytosis. One hour after phagocytosis of yeast-locked *C. albicans*, phagosomal expansion was induced by adding 200 μM GPN (left panel). HBSS was used as negative control. DIC and fluorescence images were acquired every minute for 25 min. Arrowheads point to phagosomes. Scale bar = 5 μm.

### Phagosomes undergo transient changes in pH before irreversible rupture

Biological membranes have effective means of repairing occasional tears, thereby maintaining homeostasis (39–43). It was therefore conceivable that phagosomes containing growing hyphae would attempt to maintain their integrity; at least until the repair mechanisms are exhausted. To investigate this possibility, macrophages were infected with *C. albicans* and, after hyphae were allowed to grow for 3 hrs, we repeatedly measured phagosomal pH every 5 s over a period of 10-60 min. As depicted in Figure 6A, acidic phagosomes frequently underwent a series of rapid changes in pH. We refer to this phenomenon as a ‘pH flashes’. While phagosomes often underwent one pH flash during the observation period, some phagosomes flashed up to 6 times (Movie A4). The variability of the flashing pattern is illustrated in Figure 6B, where the time courses of pH recordings from multiple phagosomes are overlaid. The rate of pH flashes was similar when the *C. albicans ece1Δ* mutant was phagocytosed, suggesting that pH flashes are a consequence of hyphal expansion rather than Candidalysin (Figure 6C and Movie A4). The flashing pattern is consistent with the notion that, during hyphal expansion, phagosomes undergo tears that can in some instances be repaired, before irreversible rupture occurs, leading to sustained alkalinization and loss of SRB. It is noteworthy that neither the transient nor the permanent phagosomal breaks were associated with macrophage lysis, as the macrophages remained largely impermeable to propidium iodide (PI) 4 hrs post-infection (Figure A3; total cell lysis was less than 10%).

**Figure 6.**
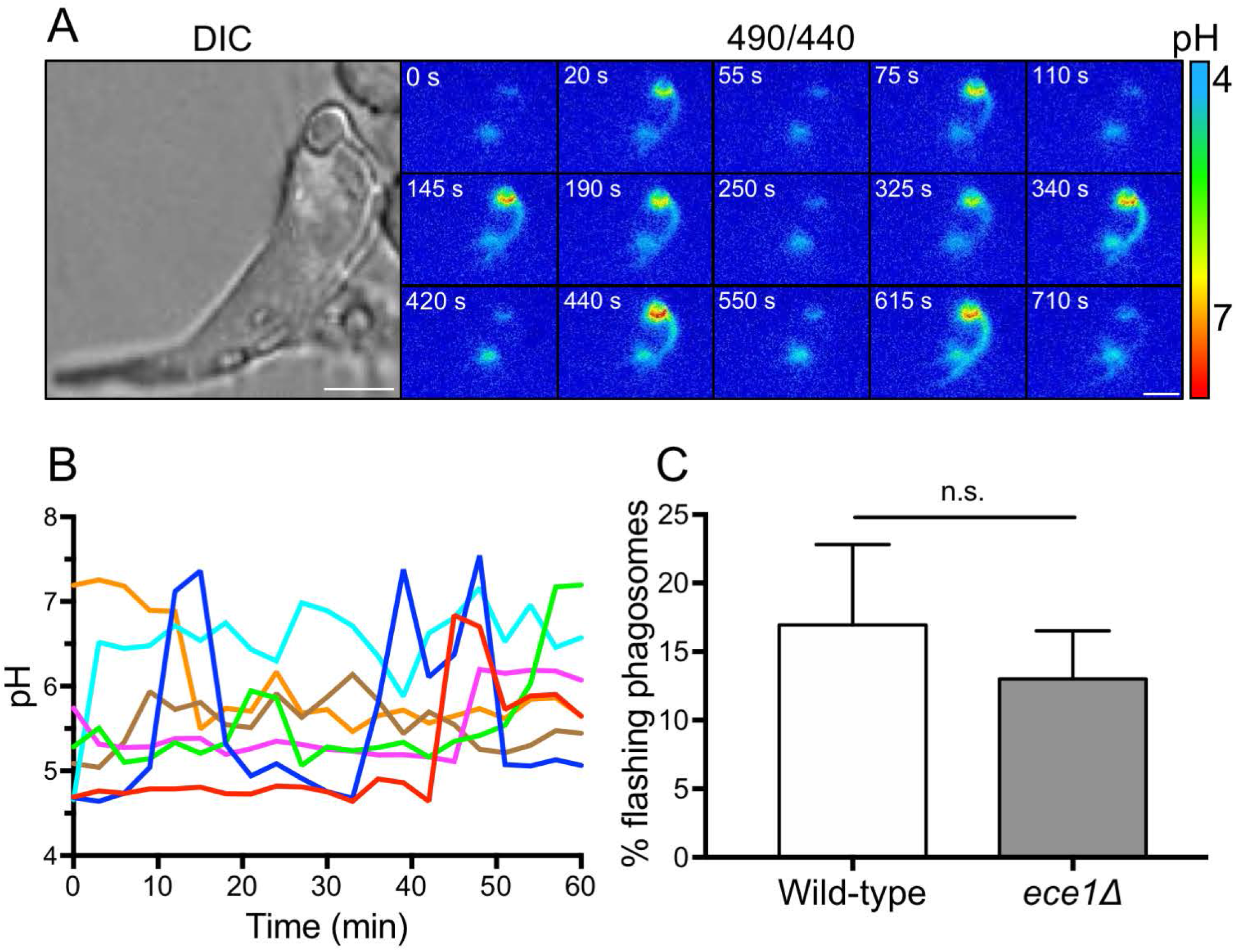
*C. albicans* phagosomes display transient proton leakage prior to undergoing irreversible breaks. (A) Phagosomal pH was measured 3 hrs post-infection by dual-wavelength ratio imaging. Images were acquired every 5 s for 15 min. An enlarged DIC image of the macrophage analyzed is shown in the left panel. Scale bars = 5 μm. (B) Different patterns of transient proton leakage recorded over 60 min. (Each individually colored traces represents a separate phagosome. C) Quantitation of the fraction of phagosomes recorded in each field of view that displayed flashing events during a 10 min period of analysis, 3 hrs post-infection. Data show means ± SEM of at least 30 determinations in 3 independent experiments of each type. Statistical significance was assessed by an unpaired t-test. n.s.= not significantly different.

## DISCUSSION

In this study, we reached three main conclusions: first, that NH_3_ generation by *C. albicans* and its retention by phagosomes cannot be responsible for the observed alkalinization; second, that initiation of hyphal growth occurs in acidic phagosomes and third, that hyphal growth drives phagosomal alkalinization by stretching and eventually rupturing the phagosomal membrane. These conclusions are discussed in turn below.

It is generally agreed that the mature phagosome containing *C. albicans* is highly acidic, and low pH is known to inhibit yeast-to-hypha transition. Hence, it was recently suggested that *C. albicans* produces NH_3_ to alkalinize the phagosome prior to hyphal growth (13, 15–18). Our findings suggest that intracellular NH_3_ production by *C. albicans* is not directly responsible for the alkalinization. The rate of NH_3_ generation is much too low to overcome the ability of the V-ATPase to acidify the phagosome (Figure 1) and, more importantly, NH_3_ cannot be retained within the phagosomes, which are extremely permeable to the uncharged weak base (Figure 2).

Our findings indicate that, rather than being the cause of hyphal growth, phagosomal alkalinization is its consequence. How then is hyphal extension initiated? Several environmental factors influence yeast-to-hypha transition, including high temperature (37°C), adherence to host cells, high CO_2_ and nutrient deprivation (4, 5). All of these factors are experienced by *C. albicans* cells within the mammalian phagosome and may suffice to induce hyphal formation despite the acidic pH, albeit at a reduced rate. Accordingly, we find that *C. albicans* hyphae grow faster in phagosomes treated with concanamycin A, a V-ATPase inhibitor that dissipates the lysosomal and phagosomal acidification, which was verified using the acidotropic dye cresyl violet (Figure A2D-E). Thus, while capable of growing inside acidic phagosomes (Figure 3), *C. albicans* hyphae indeed extend more rapidly at more alkaline pH (Figure A2E).

Some of the evidence supporting the involvement of NH_3_ stemmed from experiments using mutants with defective amino acid permeases, which ostensibly lacked the substrates to generate sufficient NH_3_. One such mutant, *stp2*Δ, was found to be unable to alkalinize phagosomes (15). In the presence of amino acids, Stp2 induces the transcription of genes leading to amino acid uptake and catabolism. This in turn produces urea, which subsequently leads to the production of NH_3_ and CO_2_ by urea amidolyases. It is noteworthy, however, that Stp2 affects the expression of several genes (44, 45), and, as a result, suffers from growth defects, particularly in environments where nutrient availability is restricted, such as the phagosomal lumen. Thus, it is impossible to distinguish whether the effects caused by deletion of the gene are due to lack of NH_3_ production or to impaired growth. Other mutants such as *ahr1Δ* used to buttress the NH_3_ hypothesis suffer from similar shortcomings, which is why we opted not to include them in our analyses (14, 17, 18).

Since NH_3_ generation appeared unlikely to account for the observed alkalinization, we sought for an alternative mechanism. Our data are consistent with the notion that hyphal growth distends the phagosomal membrane, causing leakage of proton equivalents and even larger molecules like SRB. In the early stages the ruptures are transient, possibly reflecting the activation of repair mechanisms; indeed, we have preliminary evidence that re-acidification is associated with additional fusion of LAMP-positive compartments with the phagosome (data not shown). The progressive membrane tears become irreversible thereafter, judged by the impossibility to re-establish the acidic pH. Of note, the sudden and initially reversible increases in pH cannot be readily explained by NH_3_ production, which is anticipated to be continuous, producing a gradual and sustained pH change. That mechanical stretching of the phagosomal membrane is the cause of the permeability change is supported by the observation that outward hydrostatic pressure established by osmotic means–using GPN–resulted in a similar disruption of the phagosomal membrane, with dissipation of the pH gradient and leakage of SRB (Figure 5). Whether transient or more permanent, phagosomal membrane rupture exposes the fungus and its products to the cytosolic milieu. Little is known regarding the means whereby *C. albicans*-secreted effectors activate signaling pathways within the cytosol. We therefore speculate that discontinuities in the phagosomal membrane associated with hyphal growth could contribute to inflammasome activation and pyroptosis (46–48).

In conclusion, we propose that hyphal growth is initiated inside acidic phagosomes (albeit at a reduced rate) and that alkalinization results from excessive membrane distension, which either activates mechanosensitive channels and/or causes outright rupture of the phagosomal lining. The discontinuities may initially be transient, as the membrane is repaired by fusion with other organelles (likely LAMP-positive late endosomes-lysosomes), but eventually become permanent, leading to sustained alkalinization and granting the fungus access to the richer cytosolic environment.

## MATERIALS AND METHODS

### Strains and reagents

Experiments were carried out using mouse RAW 264.7 macrophages (ATCC). RAW cells were plated sparsely in 12-well tissue culture plates (Corning Inc.) and grown overnight at 37°C in an air/CO_2_ (19:1) environment in RPMI-1640 (Wisent Inc.) supplemented with 5% (vol/vol) FBS. *C. albicans* wild type strain was the prototrophic strain BWP17/CIp30 (49). Other strains used are listed in Table A2. *C. albicans* cultures were grown in YPD medium (1% yeast extract, 2% peptone, 2% dextrose) at 30°C overnight. Cultures were washed in sterile PBS and adjusted to the required cell density. Concanavalin A labeled with FITC and sulforhodamine B were from Invitrogen. Nigericin, monensin and Gly-Phe β-naphthylamide were from Sigma. Concanamycin A was from Abcam. Recombinant pneumolysin was a kind gift from Dr. John Brumell.

### Phagocytosis of *C. albicans*

After overnight incubation at 30°C, *C. albicans* yeast were washed twice in PBS and incubated in the dark with concanavalin A-FITC (1:100) and rabbit anti-*C. albicans* IgG (1:167, for 60 min at room temperature rotation). After labeling, yeast were washed twice in PBS and diluted to OD_600nm_ 1.0 in PBS.

Five μL of concanavalin A-FITC-labeled yeast in fresh RPMI-FBS were added to RAW cells grown on glass coverslips, which were then centrifuged at 1500 × g for 1 min at room temperature to synchronize phagocytosis. After 20 min incubation at 37°C, yeast not associated with the macrophages were washed away with PBS, yeast that had adhered to the macrophages but had not been internalized were labeled with donkey anti-rabbit Cy3 (1:1000) for 10 min at 37°C. Cells were imaged live or, where indicated, fixed with 4% paraformaldehyde for subsequent analysis. All phagocytosis experiments were imaged in Hank’s Balanced Salt solution (HBSS).

### Buffering capacity (β)

To determine buffering capacity (β) of the Candida-containing phagosome, phagosomes were allowed to acidify for 1 h before phagosomal pH was measured ratiometrically, as described below. The bathing solution was switched to HBSS containing 15 mM NH_4_Cl, and the pH was measured again immediately. At the end of each experiment, a standard calibration was performed as described below and fluorescence ratios were converted to pH. The intra-phagosomal NH_4_^+^ concentration ([NH_4_^+^]) was calculated using the Henderson-Hasselbalch equation, and the intrinsic buffer capacity (in millimoles/liter/pH) was calculated as Δ[NH_4_^+^]/ΔpH.

### Vacuolar H+ ATPase (V-ATPase) pumping rate

Phagosomes were generated as described and, after 1 h, the steady-state phagosomal pH was measured prior to addition of 2 μM CCA to the bathing solution. Thereafter, the phagosomal pH was measured every minute for 15 min at 37°C. At the end of each experiment, a standard calibration was performed and fluorescence values converted to pH. The rate of change of the luminal pH (ΔpH) measured during the first minute after CCA treatment was used to estimate the V-ATPase pumping rate, which was assumed to be identical to the proton leakage rate at steady state. Proton pumping rates were calculated as (ΔpH * β * phagosome volume) / time. To quantify the phagosomal volume, the radius of phagosomes containing *C. albicans* yeast was measured microscopically. Volume was calculated assuming that the phagosomes were spherical (volume = 4/3 π r^3^). Phagosomal and lysosomal alkalinization was confirmed using the acidotropic dye cresyl violet as previously described (50).

### Determination of NH_3_ leakage rate

Phagosomes were generated as described and, after 1 h, the steady-state phagosomal pH was measured prior to the addition of 15 mM NH_4_Cl. Phagosomal pH was then measured every second for 10 seconds. A standard calibration was generated as described below, to determine the change in pH (ΔpH) induced by NH3 addition or withdrawal. Leakage of NH_3_ was calculated as (ΔpH * β * phagosome volume) / time.

### Dual-wavelength ratiometric fluorescence measurements

RAW cells were allowed to ingest *C. albicans* as described above and the coverslips were then mounted in a Chamlide magnetic chamber and overlaid with HBSS. The chamber was placed in a Leiden microincubator maintained at 37°C on the stage of an inverted microscope (DM IRB; Leica Biosystems) equipped with a 40A/1.25 N.A. oil objective (Leica Biosystems), a lamp (X-Cite 120; EXFO Life Sciences Group), and filter wheels (Sutter Instrument) that control excitation and emission filters. For experiments using concanavalin A-FITC-labeled *C. albicans*, excitation wavelengths were alternated between 485 ± 10 nm and 438 ± 12, with emitted light selected through a 520 nm filter. Light was captured by a cooled electron-multiplied charge-coupled device camera (Cascade II; Photometrics). The filter wheel and camera were under control of MetaFluor software (Molecular Devices). Emission at 520 nm from light excited at the two excitation wavelengths was collected and their ratio was calculated on-line from phagosomes at different time points. For phagosomal pH measurements, the 490/440 nm ratio of at least 30 phagosomes was measured for each strain and time point. For transient pH oscillations (pH flashes), the 490/440 ratio was acquired from the same field of view every 2 sec for 10 min, or every 3 min for 60 min. For GPN-induced phagosomal expansion, 100 μM GPN was added to phagosomes containing yeast-locked *C. albicans* 1 h post-infection. GPN-treated phagosomes were visualized for 30 min at 37°C.

### Conversion of dual-wavelength fluorescence ratio to pH

To convert measured phagosomal fluorescence ratios to pH, samples were sequentially bathed in isotonic K+ solutions containing 10 μM nigericin and 5 μM monensin, and calibrated to pH 7.5, 6.0, 5.5, 5.0 and 4.5 respectively. Samples were imaged 5 min after addition of each solution to ensure pH equilibration across all compartments. After background subtraction at each wavelength, measured fluorescence ratios at defined pH were plotted into a calibration curve that was fit with least squares. The measured phagosome fluorescence ratios were transformed into intracellular pH by using the equation describing the curve generated above. For pH measurements, fluorescence values from each phagosome were obtained using the freehand tool in Fiji (version 1.0) to select regions of interest (ROIs) delimiting the phagosome. The fluorescence intensity was corrected by subtracting the background fluorescence at each wavelength and converted to pH by using the equation described above. For pseudocolor display, a RatioPlus plug-in in Fiji was used to depict the 490/440 ratio.

### *C. albicans* hyphal growth

The length of *C. albicans* was measured from DIC images acquired at different time points after phagocytosis. Total length included the yeast head and hyphal projection.

### Sulforhodamine B loading and leakage during phagocytosis

To load endosomes and lysosomes with SRB, RAW cells were bathed for 60 min at 37°C in RPMI-FBS containing 150 μg SRB/mL, prior to phagocytosis. After loading, phagocytosis was initiated as described above. Leakage of SRB from phagosomes was measured using spinning disk confocal microscopy, and the percentage of SRB-positive phagosomes was calculated as (SRB-positive phagosomes/total phagosomes) * 100. For GPN-induced membrane rupture, 200 μM GPN was added to phagosomes containing yeast-locked *C. albicans* 1 h postinfection. GPN-treated phagosomes were visualized for 30 min by DIC and fluorescence microscopy.

### Spinning disk confocal microscopy

Confocal images were acquired using a spinning disk system (WaveFX; Quorum Technologies Inc.). The instrument consists of a microscope (Axiovert 200M; ZEISS), scanning unit (CSU10; Yokogawa Electric Corporation), electron-multiplied charge-coupled device (C9100-13; Hamamatsu Photonics), five-line (405, 443, 491, 561, and 655 nm) laser module (Spectral Applied Research), and filter wheel (MAC5000; Ludl) and is operated by Volocity software version 4.3.2 or 6.2.1 (Perkin-Elmer). Images were acquired using a 63x/1.4 NA oil objective (Zeiss) coupled to an additional 1.5x magnifying lens and the appropriate emission filter. Cells were maintained at 37°C using an environmental chamber (Live Cell Instruments).

### Statistical analysis

Unless otherwise indicated, data are presented as means ± SEM of the number of determinations shown in parenthesis. Statistical significance was determined using unpaired t-test, one-way ANOVA (Tukey’s test or Dunnett’s test) and two-way ANOVA (multiple comparisons) with PRISM 7 (GraphPad Software), with p< 0.05 considered significant.

## ACKNOWLEDGEMENTS

We thank Thomas Krüger and Olaf Kniemayer (MAM, HKI) for LC/MS support.

